# Beyond Threat: Changes in Visuocortical Engagement and Oscillatory Brain Activity during Non-Aversive Associative Learning

**DOI:** 10.1101/2025.07.14.664451

**Authors:** S. M. Gardy, C. Panitz, H Engle, F. Gilbert, A. Keil

**Author notes:** Corresponding Author: Sarah M. Gardy, Department of Psychology Laboratory for Brain, Body, and Behavior, University of Florida.

## Abstract

Aversive conditioning produces selectively heightened visuocortical responses to conditioned stimuli stimulus (CS) that predict aversive unconditioned events (US). However, it is unclear whether similar neural signatures emerge as a consequence of mere association formation between a CS-US pair, i.e., when a CS predicts a neutral event. To address this, we paired a soft tone (65 dB) with one of two high-contrast circular gratings (15° or 75°; CS+; counterbalanced), while two intermediate orientations (35°, 55°) were never paired with the neutral tone, serving as generalization stimuli (GS). A sample of 22 participants viewed each grating for 3000 ms, with the tone presented during the last 1000 ms of each grating presentation. Gratings were flickered (turned on and off) at a temporal rate of 15 Hz, to evoked steady-state Visual Evoked Potentials (ssVEPs), a metric of visuocortical engagement. Time-frequency decomposition via Morlet wavelets quantified changes in the amplitude of alpha-band (8-13 Hz) oscillations—an additional electrophysiological index that has been shown to be sensitive to aversive conditioning. Contrary to findings observed during aversive conditioning, alpha amplitude increased, rather than decreased, during CS+ trials relative to GSs. Likewise, ssVEP amplitude was higher for GSs than for the CS+, which again is the opposite of what is found during aversive conditioning. These findings suggest that a CS paired with non-aversive outcomes engages mechanisms consistent with working memory, anticipation, or imagery processes, reflected in heightened alpha amplitude and attenuated ssVEP, rather than the defensive potentiation observed during aversive conditioning.

## Introduction

Conditioning is a fundamental process by which organisms form associations between stimuli and outcomes. This associative process supports adaptive behavior by allowing individuals to anticipate and respond to significant environmental events (Pavlov, 1927). Much of the research in this field has focused on how associations form between aversive and neutral stimuli, often referred to as aversive conditioning (Hamm & Weike, 2005). This research has also had strong translational implications, informing, for example, understanding of anxiety and trauma-related disorders (Craske et al., 2008). Aversive conditioning experiments involve pairing a neutral stimulus (CS+) with an aversive unconditioned stimulus (US). Eventually, the CS+ becomes predictive of the US and reliably elicits an aversive response. Other cues (the CS-) are never paired with the US and thus do not elicit aversive responses. The ability to rapidly acquire aversive associations is thought to be evolutionarily advantageous, providing a protective mechanism against acute threats (Lang & Bradley, 2010).

A substantial body of research exists examining aversive learning paradigms and the impacts of conditioning on neurophysiology, particularly in the visual domain. Studies using steady-state visually evoked potentials (ssVEPs), oscillatory brain responses that are phase-locked to a flickering visual stimulus, have shown increased amplitude to threat-predictive stimuli (Li & Keil, 2023; Miskovic & Keil, 2015; Panitz et al., 2019; Song & Keil, 2014). These responses, primarily generated in early visual cortex (Mueller et al., 1997), reflect heightened sensory gain and are interpreted as indicators of enhanced visuocortical engagement with a given stimulus. Several studies have examined how this heightened visuocortical engagement generalizes to cues that vary in similarity to the CS+. Generalization conditioning paradigms are used towards this goal. In these paradigms, stimuli systematically differing in similarity to the CS+ are not paired with an aversive stimulus. Differential conditioning paradigms using different feature-similarity gradients have shown that ssVEPs become finely tuned to the CS+, supporting precise discrimination between threat and safety cues (Ahumada et al., 2025; McTeague et al., 2015). Additionally, research has demonstrated that ssVEPs become amplified in the presence of other aversive cues such as pictures (Keil et al., 2005), in occipital and parietal areas (Keil et al., 2009).

Complementing ssVEP findings, alpha-band oscillations (8-13 Hz)—the dominant EEG rhythm over occipital and parietal regions—have been hypothesized to index cortical excitability and attentional processing (Klimesch, 2012). Often thought to reflect activity in thalamo-cortical circuits (Başar et al., 1997), changes in the magnitude of alpha oscillations have been connected with processing of external stimuli and have been hypothesized to have an inhibitory role in brain function (Klimesch, 2012). In many current electrophysiological models, alpha amplitude is inversely related to cortical excitability, where higher alpha amplitude is associated with reduced sensory processing and increased inhibition (Jensen & Mazaheri, 2010). By contrast, lower alpha amplitude is thought to reflect heightened neural excitability and improved perceptual accuracy (Friedl & Keil, 2021). During aversive conditioning, CS+ have been shown to elicit greater suppression of alpha-band amplitude compared to safety cues when using simple visual stimuli (Bacigalupo & Luck, 2022; Panitz et al., 2019) as well as when using more complex conditioned stimuli such as faces (Pouliot et al., 2024). This alpha suppression is interpreted as a neural signature of increased attentional engagement with the threat-predictive cue.

While aversive conditioning research has informed our understanding of association formation, it is unclear whether the observed effects of aversive conditioning on visuocortical responses are specific to defensive/aversive association formation or if they reflect a broader mechanism of association formation. Most existing studies have focused exclusively on emotionally salient stimuli, leaving open the question of how early sensory cortical activity, and visual attention overall, is modulated by the formation of non-aversive associations. Evidence from animal models suggests that associative plasticity is not restricted to aversive contexts. For example, Headley and Weinberger (2015) found that after repeated pairings of neutral light and tone stimuli, neurons in the rat primary visual cortex (V1) developed responses to the auditory tone. This effect was not observed in control groups exposed to unpaired stimuli, indicating that this type of associative plasticity is learning dependent, but does not require aversive US.

### The Present Study

Despite extensive research on aversive conditioning, studies examining neurophysiological associative learning responses in the absence of emotionally or motivationally salient stimuli are scarce. The present study seeks to fill this gap by adapting an established aversive conditioning paradigm (Ahumada et al., 2024), substituting the aversive white noise with a soft tone (65 dB). Using ssVEPs and alpha amplitude as indices of visuocortical engagement, we investigated whether associative learning between neutral visual and auditory stimuli elicit comparable patterns of sensory modulation in the absence of aversive stimuli. This work contributes to a broader understanding of the neural basis of associative memory formation and helps to delineate activity specific to emotionally salient learning from those reflecting general associative processes.

## Methods

A total of 22 participants were included in the study (17 female). This sample size mirrors the original studies in which effects of aversive conditioning on ssVEPs and on alpha amplitude changes were observed (e.g., McTeague et al. 2015). Participants ages ranged from 18-23, with an average age of 19.6. Gender distribution included 17 women and 5 men. Racial breakdown of participants included 16 white, 2 black, 3 Asian, and 1 other, with 5 identifying as Hispanic. Exclusion criteria stipulated that participants be at least 18 years of age, have normal or corrected normal vision, and have no history of epilepsy. Individuals gave written consent to participant in the study and received class credit in compensation for their time. The experiment was conducted in accordance with the Declaration of Helsinki and approved by the University of Florida Institutional Review Board.

## Materials and Procedure

Participants viewed four black-and-white, high-contrast (Michelson contrast 100%), circular Gabor gratings with variation in orientation. A soft tone of 65 dBA, measured with a sound meter located at the ear position of the participants, serving as US, was paired with either a 15° or 75° grating (CS+, counterbalanced across participants), while 35° and 55° orientations were never paired with the tone (Generalization stimuli, GSs). The gratings were presented for 3000ms, flickered at a rate of 15 Hz to elicit ssVEPs. The soft tone was played during the last 1000 ms of the CS+, co-terminating with the grating. Analyses were conducted on the first 2000ms of grating presentation (i.e., the timespan before tone onset).

**Figure 1.**
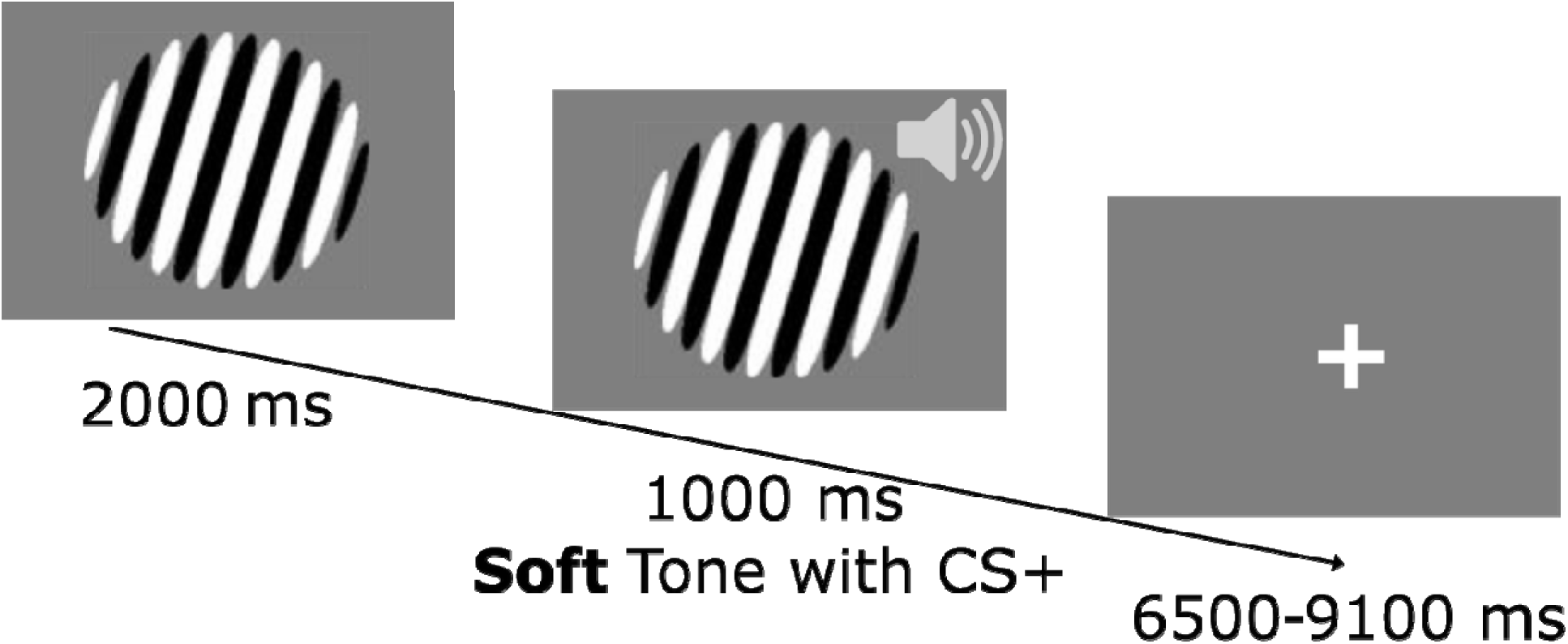
Trial structure illustration with CS+, where Gabor is presented for 2000 ms and soft tone occurs for 1000 ms. Co-terminate for an inter-trial-interval between 6500-9100 ms. Gabor stimuli flickered at 15 Hz.

Stimuli were presented using MATLAB Psychophysics Toolbox code (Brainard, 1997; Pelli, 1997) on a Cambridge Research Systems Display ++ monitor (1,920 x 1,080 pixels, 120 Hz refresh rate) at 5° of visual angle in the center of a medium gray (RGB: 127, 127, 127; 45 cd/m^2^) background. The soft tone was delivered through two studio monitor speakers (Behringer Studio 50) located directly behind the participant. Participants were seated approximately 125 cm from the screen.

After the researcher explained the study, participants reviewed and signed the consent form. They were then guided to the experimental chamber and completed questionnaires on trait anxiety, social anxiety, depression (not discussed in this manuscript). Participants were seated in front of the screen, instructed to maintain fixation in the middle of the screen and minimize movement during the task, and were fitted with the EEG net.

The Pavlovian conditioning paradigm consisted of a habituation, acquisition, and extinction phase. During the habituation phase, participants viewed each grating 8 times, for a total of 32 trials, and no tone was presented. Valence, arousal, and expectancy ratings were collected at the end of this phase. During acquisition, CS+ gratings (either 15° or 75°) co-terminated with the tone US. The first 6 CS+ trials co-terminated with the tone US 100% of the time, followed by 50% contingency for the 18 remaining trials. The GS gratings were each presented 24 times for a total of 96 presentations. Acquisition was divided into two blocks with ratings taken in the middle and at the end of the phase. Extinction consisted of 12 trial presentations for each of the 4 gratings for a total of 48 trials, with ratings taken again at the end. The inter-trial interval ranged from 6500-9100 ms. Participants were not informed of the contingencies between the CS+ and the US.

Valence and arousal were measured on a scale of 1 to 9, with 1 indicating more unpleasantness and low arousal, and 9 indicating more pleasantness and high arousal, respectively. Expectancy was measured as the participants’ perceived probability of the US occurring given a certain grating.

### Data Recording and Processing

#### EEG

EEG was recorded with 128-channel HydroCel Geodesic sensor nets and a Net Amp 300 amplifier (Magstim EGI, Oregon, US). The sample rate was 500 Hz and referenced to the Cz electrode. On online Butterworth filter was set at 200 Hz. Electrode impedances were kept below 60 KOhms when feasible.

For EEG preprocessing, a version of the SCADS (Statistical Correction of Artifacts in Dense Array Studies) method (Junghöfer et al., 1997) was used for data preprocessing in MATLAB (version 2022, Natick, MA, USA). The continuous signal was filtered using a 4^th^ order Butterworth low-pass filter with a 3dB point at 30 Hz and a 4^th^ order high-pass filter with a 3 dB point at 1 Hz. Continuous data were segmented into 3.6 second epochs (1801 sample points), spanning 600 ms before cue onset to 3000 ms after tone onset. Segmented trials were evaluated for artifact contamination using SCADS which utilizes mean amplitude, standard deviation, and gradient voltage amplitude. Channels were flagged as globally bad if data artifacts exceeded 2.5 standard deviations above a median quality index. The globally bad channels were interpolated with spherical spline interpolation (Junghöfer et al., 1997). For each trial, a quality index threshold of 2.5 standard deviations was used, where trials were flagged and interpolated within trial with 2D spline interpolation. Eye artifacts were corrected with a regression-based EOG method (Schlögl et al., 2007; 2009).

#### SSVEP

SsVEP amplitudes were calculated via two methods. First, to isolate the ssVEP at 15 Hz and maximize its signal-to-noise ratio, we applied the rhythmic entrainment source separation (RESS; Cohen & Gulbinaite, 2017) procedure on each participant’s artifact-free trials, in a time window of 2000 ms between the Gabor onset and the onset of the soft tone. This method uses electrode channels across time and trials to find an optimal spatial filter for the ssVEP driving frequency while minimizing the filter’s sensitivity to neighboring frequencies. Spatial weights are derived from an eigenvalue decomposition to optimally separate the driving frequency from the neighboring frequencies. The resulting RESS time series for each trial were transformed into the frequency domain by means of Discrete Fourier Transform (DFT), with a cosine-square of 20 points duration applied at the beginning and at the end, to minimize leaking and spectral distortion. The resulting power spectrum was obtained as the absolute value of the sine and cosine components, and normalized by the number of points entering the DFT. These single-trial spectra were then averaged within participant and condition. The power at 15 Hz was measured from the resulting participant by condition power spectra and used for subsequent analyses.

The second method for determining the ssVEP amplitude at 15 Hz mirrored the approach used in Ahumada et al. (2024). We applied a DFT as described above directly to each artifact-free single trial to compute per-channel amplitude spectra over the same 2000 ms time window as described above for the RESS method. The resulting single-trial spectra were then averaged within participant, condition, and electrode, yielding a participant by condition by electrode matrix containing 15 Hz ssVEP amplitude, used for subsequent analyses. This additional approach was employed as an added control, and to examine the generalizability of findings beyond the RESS method.

### Alpha amplitude

A Morlet wavelet analysis quantified alpha-band activity, and spectral estimates of the ssVEP amplitude were calculated. All statistical analyses were computed with MATLAB (version 2023, Natick, MA, USA). Artifact-free single trials were transformed into the time-frequency domain using a family of Morlet wavelets with a constant Morlet parameter m=10. Wavelets were computed as the convolution of the wavelets with the data for frequencies between 2.8 and+ 29.4 Hz, in steps of 1.11 Hz. These settings resulted in a frequency uncertainty (sigma_f_) of 1.055 Hz and a time uncertainty (sigma_t_) of 151 ms at the frequency of 10.56 Hz, chosen for analysis. The single-trial time-frequency data were averaged together and baseline adjusted via subtraction from 200 ms prior to stimulus onset. An average of 6.27% of channels were interpolated for each participant with an average of 14.36 % trials removed per participant. An average of 99.64 trials remained for CS+, 99.55 for GS1, 99.14 for GS2, and 100.05 for GS3. As stated above, the alpha frequency examined here was operationalized as the amplitude of the wavelet corresponding to 10.56 Hz.

### Statistical Analyses

The goal of the present paper was to examine association formation in a non-aversive conditioning paradigm. To this end, we first examined rating differences across condition and phase for arousal, valence, and expectancy. Bayesian repeated measures ANOVAs were conducted on all three sets of ratings with Cue (CS+, GS1, GS2, GS3) and Phase (Habituation, Acquisition 1, Acquisition 2, Extinction) as within-subjects factors, implemented in JASP (JASP Team, 2024). Prior model probabilities for each rating questionnaire were set as equal (P(M) = .20).

Next, we evaluated dependent variables using the Bayesian Bootstrap approach (Efron, 2011) with a-priori models represented by weight vectors (Figure 2). This approach directly tests the generalization functions without requiring many individual comparisons between the conditions. We and others have used this approach before in studies of generalization conditioning (i.e., Stegmann et al., 2020; Ahumada et al., 2024). The generalization model represents a graded response across stimulus categories with alpha weights of (−2, −1, 1, 2) and ssVEP weights of (2, 1, −1, −2) applied to the CS+, GS1, GS2, and GS3, respectively. The all-or-nothing model captures a selective response to the conditioned stimulus relative to the GSs with alpha weights of (−3, 1, 1, 1) and ssVEP weighs of (3, −1, −1, −1). Note that these a-priori models are not orthogonal and are not mutually exclusive but reflect what has been found in several studies with aversive US. Weights were chosen so that they sum to zero.

**Figure 2.**
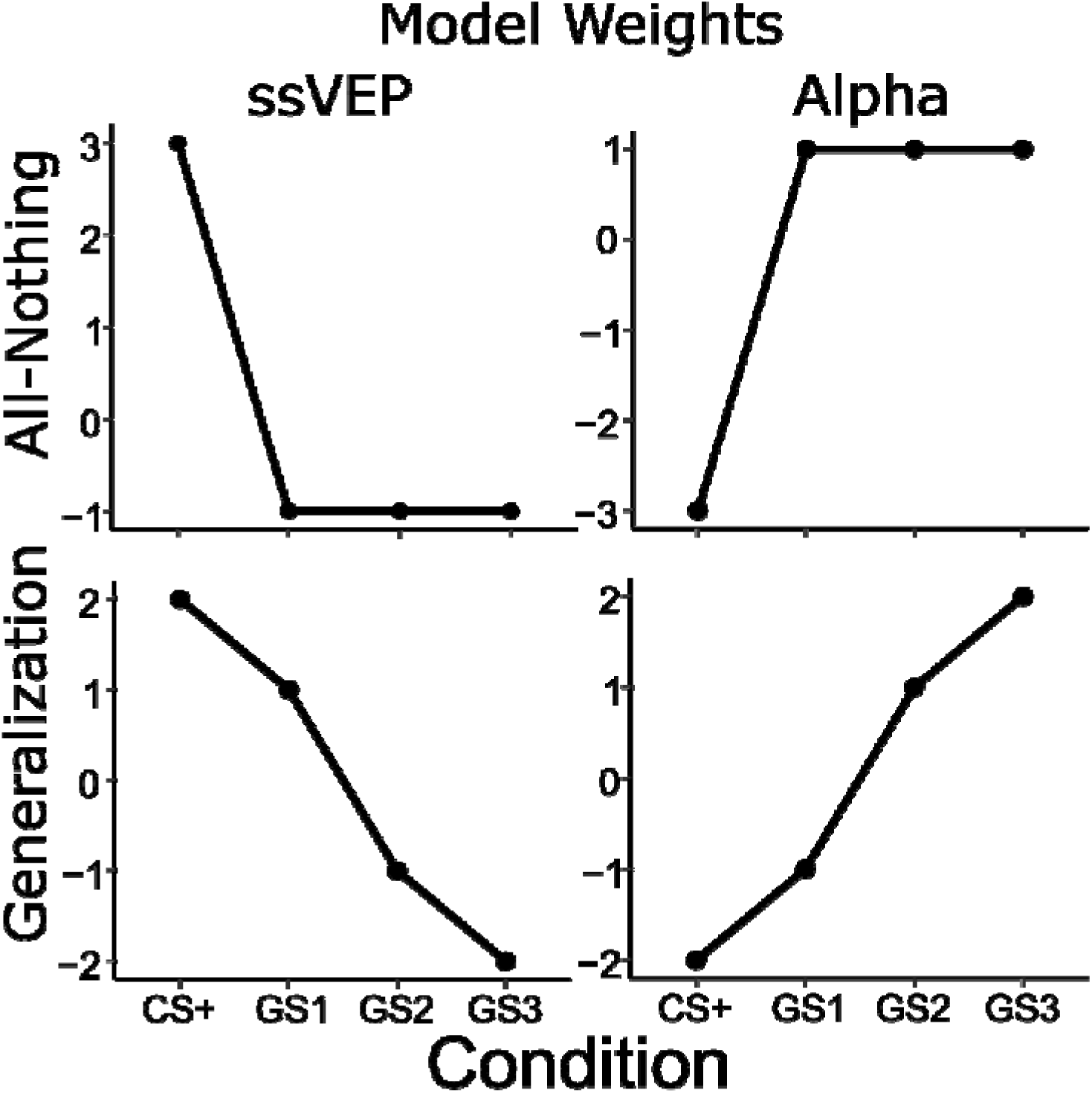
A priori pattern weights that are hypothesized to emerge during associative learning. Right column: weights for ssVEP activity. Left column: weights for alpha activity. Top row: all-nothing weights for both responses of interest. Bottom row: generalization weights for both responses of interest.

Analyses were conducted on the first 2000 ms of grating presentation, which corresponds to the timespan after Gabor onset and before soft tone onset. For transient alpha, time was divided into three epochs, Early (0-700 ms), Middle (700-1300 ms), and Late (1300:2000 ms). Bootstrapped Bayes Factors were used for analyses where posterior distributions were estimated by means of bootstrapping the data for each model and comparing the distributions to a null model (Efron, 2011).

Each of the three methods used the same three-stage statistical approach for determining Bayes Factors in favor of the models, with variation only in the dimensions of the input data and the resulting matrix. For the RESS method, at each electrode and frequency bin, we generated a bootstrap distribution of the inner product between data and the a-priori models (generalization, all-or-nothing), effected by repeatedly (2000 times) resampling subjects with replacement, computing the mean response across participants in each condition, and computing the inner product between the resulting vector with each of the model weights. This yields an empirical posterior distribution of the effect estimate under the respective a-priori model.

To approximate the null distribution, we first permuted conditions within each subject and then repeated the same resampling and bootstrapping procedure 2000 times. This preserves within-subject variance. Finally, we compared the observed posterior distribution versus the null contrast distribution at each electrode-frequency sample using the following procedure. First, the distributions were jointly z-scored to preserve differences in mean and variability, while the combined distribution has a mean of 0 and a SD of 1. Each z-scored vector was fit with a Gaussian distribution via maximum-likelihood estimation. We evaluated each fitted normal probability density function and approximated the probability mass above zero by summing the densities at positive grid points (divided by step size). From these values we computed posterior odds

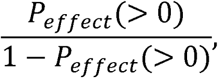

and prior odds

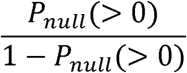

Then formed the Bayes Factors as their ratio. Bayes Factors were converted to the log 10 scale, and following convention, log_10_(BF_10_) > 1 was considered strong evidence for a model trend and log_10_(BF_10_) < −1 as strong evidence for an opposite trend (Jeffreys, 1998). All subsequent thresholding and plotting were performed on the log-transformed scale.

For the single spectral ssVEP analysis, we applied an identical Bayesian Bootstrap pipeline on a 129-electrode x 500 frequency bin x 22 subject x 4 condition data matrix. Similarly, for the alpha amplitude analysis we applied the same three-stage pipeline but applied to a 129-electrode x timepoint x 500 frequency bin x 22 subject x 4 condition data matrix. Alpha data were downsampled to 180 sample points to allow for faster computation. Alpha amplitude data were grouped by condition for analyses on condition differences.

## Results

### Ratings

Ratings for valence, arousal, and expectancy are presented in Figure 3A-C. Results revealed that valence ratings remained stable across both condition and phase, indicating that participants did not view stimuli as any more or less pleasant or unpleasant over the course of the task.

**Figure 3.**
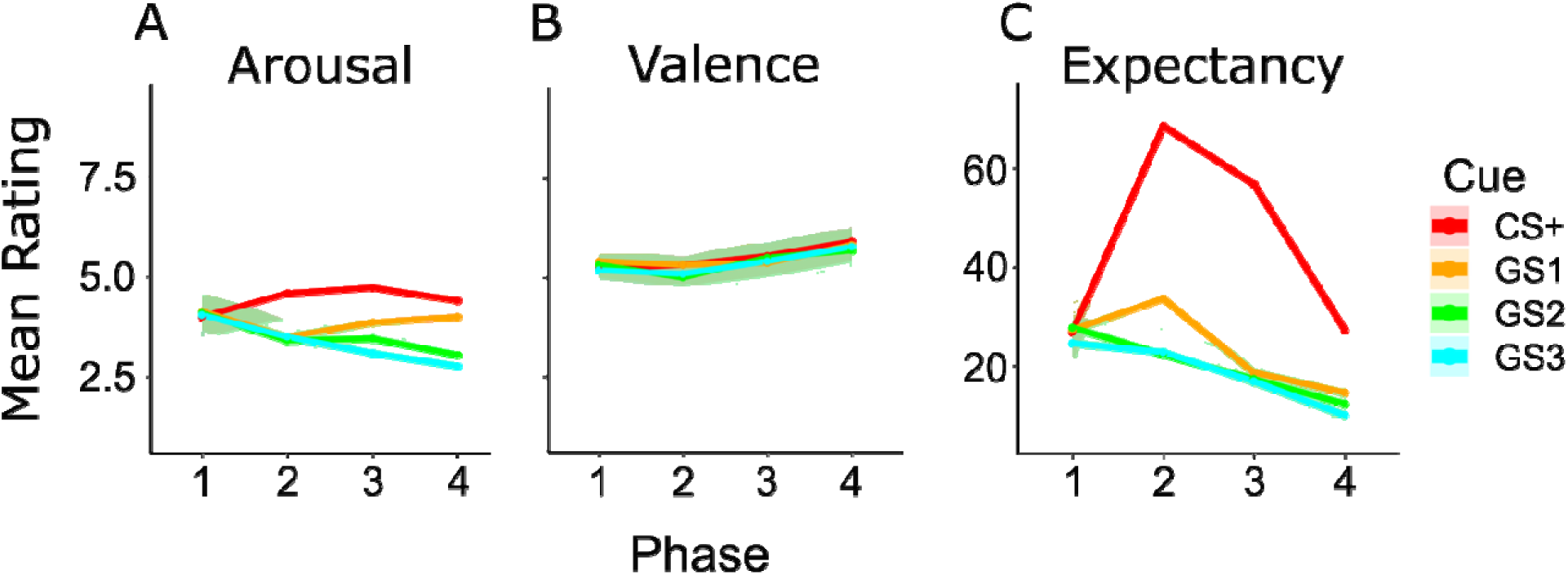
Ratings averaged over all participants by cue (CS+, GS1, GS2, and GS3) and phase (1 = Habituation, 2 = Acquisition 1, 3 = Acquisition 2, 4 = Extinction). Arousal and Valence are shown on the same scale. A) Arousal ratings. B) Valence ratings. C) Expectancy ratings.

For arousal, a cue-only model was strongly favored with a posterior model probability of P(M|data) = .67 and a Bayes Factor of BF_10_ = 61.7, indicating strong evidence for a cue effect relative to the null model. Adding Phase did not improve the fit: the Cue + Phase + Interaction model was less likely (*P*(M | data) = .23; BF_10_ = 21.5) and the Cue + Phase model had weaker evidence (BF_10_ = 7.7). The phase-only model was decisively disfavored (BF_10_ = .13). Post hoc contrasts for condition showed decisive evidence that the CS+ cue elicited greater arousal ratings than both the GS2 (BF_10_ = 422) and GS3 (BF_10_ = 970), and moderate evidence for a greater rating response compared to GS1 (BF_10_ = 5.80). Additionally, the three GSs did not differ meaningfully from one another: evidence was weak for GS1 – GS3 (BF_10_ = 1.28), GS1 – GS2 (BF_10_ = 0.75), and GS2 – GS3 (BF_10_ = 0.21). Taken together these results support the idea that elevated arousal ratings are specific to the CS+, whereas the GSs do not differ from each other.

For valence, the null model had the highest posterior probability, (P(M|data) = .69). The phase only model was the next best (*P*(M | data) = .27), yet the Bayes factor relative to the null was BF_10_ = 0.39, indicating that the data were about 2.6 times more likely under the null than under a phase effect. Models that included cue, either alone (BF_10_ = 0.04) or combined with Phase (BF_10_ = 0.017), were not supported (BF_10_ = 1.67 x 10^-4^). Findings indicate that valence ratings did not vary systematically with condition or learning phase.

Finally, the models containing main effects of cue, phase and their interaction was overwhelmingly preferred (P(M|data) = 1.00, BF_10_ = 9.94 x 10^23^), indicating that expectancy varied with phase and with condition. Post-hoc contrasts for phase revealed that expectancy increased from habituation to early acquisition (BF_10_ = 5.55) and again from early to later acquisition (BF_10_ = 2.84 x 10^3^), then dropped by extinction (BF_10_ = 5.06 x 10^3^). Condition contrasts show that CS+ consistently evoked higher expectancy than the GSs: CS+ > GS3 (BF_10_ = 2.95 × 10), CS+ > GS2 (BF_10_ = 5.37 × 10), and CS+ > GS1 (BF_10_ = 8.13 × 10). Among the GS conditions, there was no evidence for a difference between GS2 and GS3, (BF_10_ = 0.31), or GS1 and GS2 (BF_10_ = 0.73), though GS1 and GS3 were moderately different (BF_10_ = 4.27). These findings demonstrate that expectancy ratings are modulated by condition and learning stage, where it is highest for acquisition phases, declines with extinction, and is consistently greatest for the CS+ relative to the GSs.

### Steady-State Visually Evoked Potentials

Figure 4 shows the raw ssVEP data in the time (1^st^ panel) and frequency (2^nd^ panel) domains.

**Figure 4.**
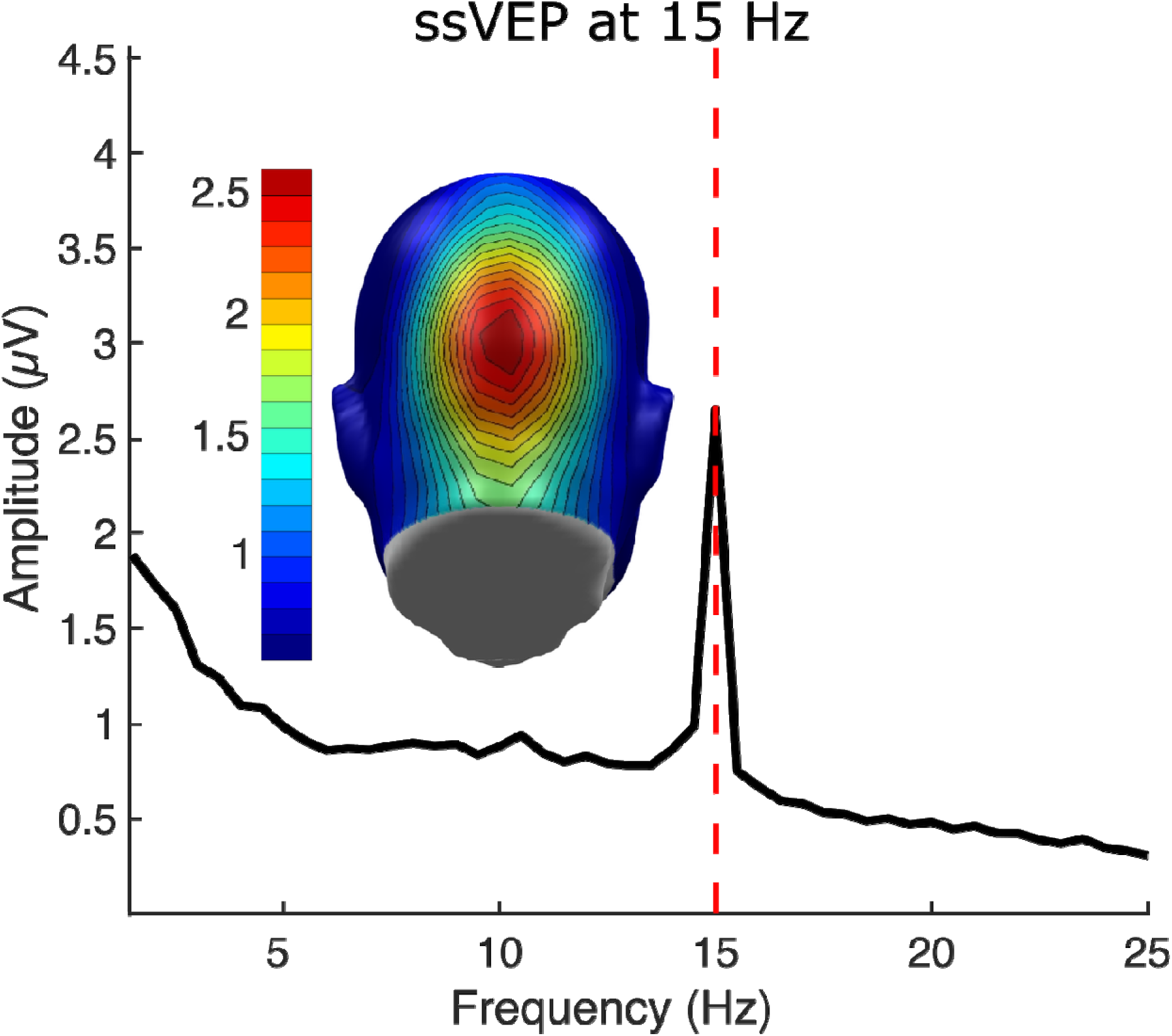
ssVEP data. A) Time domain recorded from channel Oz, averaged across all GS conditions and participants from before cue onset to co-termination of the cue and tone. B) Frequency domain displaying 15 Hz across channels in head topography and spectrum of frequency amplitudes.

Two methods were used to examine ssVEP results. The first was the RESS method, the spatial filtering method designed to maximize signal-to-noise ratio of steady-state responses by separating neural activity at the stimulus frequency from neighboring frequencies. Bayes factor comparisons show no evidence in favor of a generalization pattern at 15 Hz (log_10_BF_10_= −0.52) or an all-or-nothing pattern (log_10_BF_10_ = -.32). See Figure 5.

**Figure 5.**
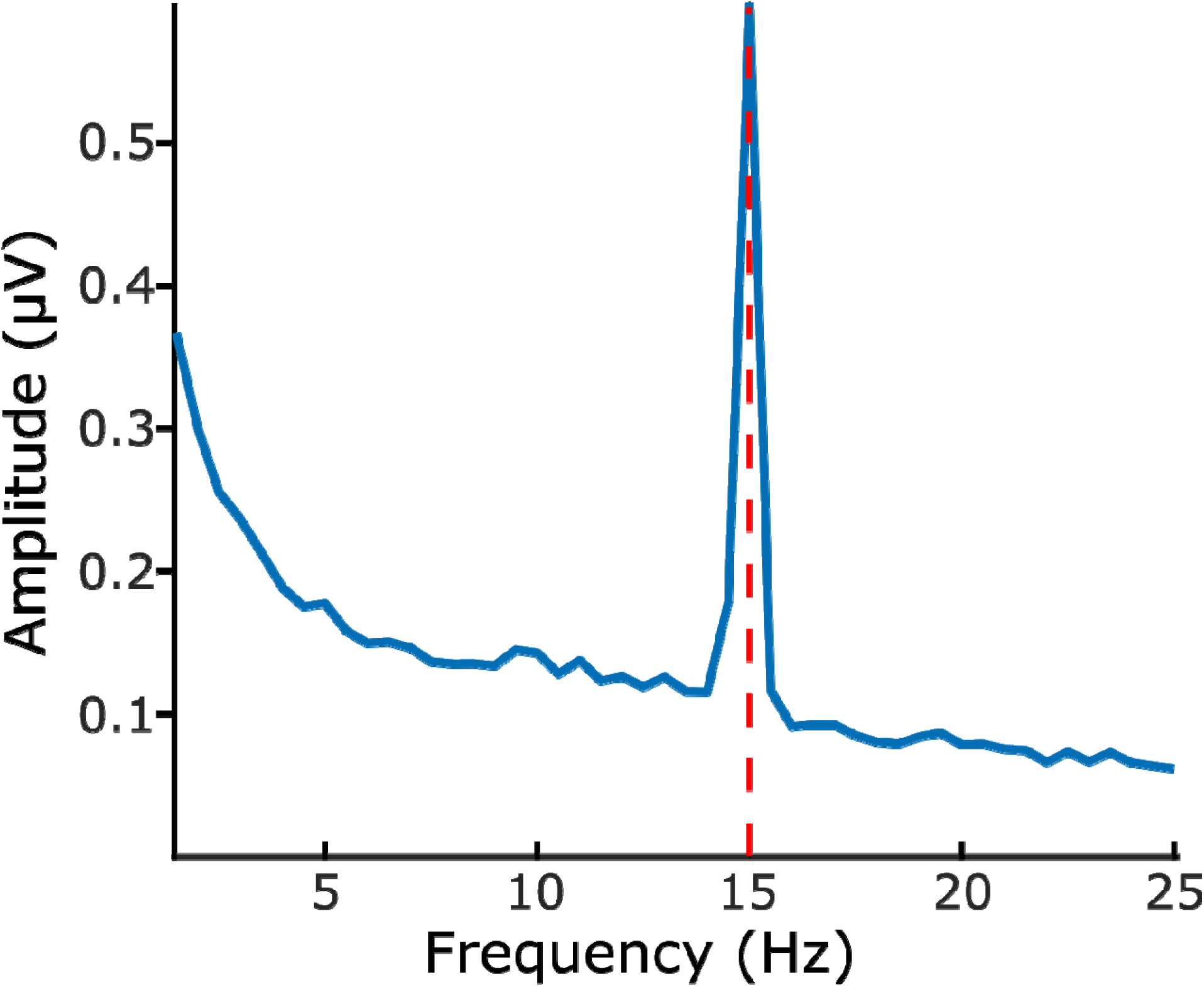
RESS filtered frequency domain displaying ssVEP activity at 15 Hz. Amplitude is lower across frequencies than for non-RESS filtered spectra.

**Figure 6.**
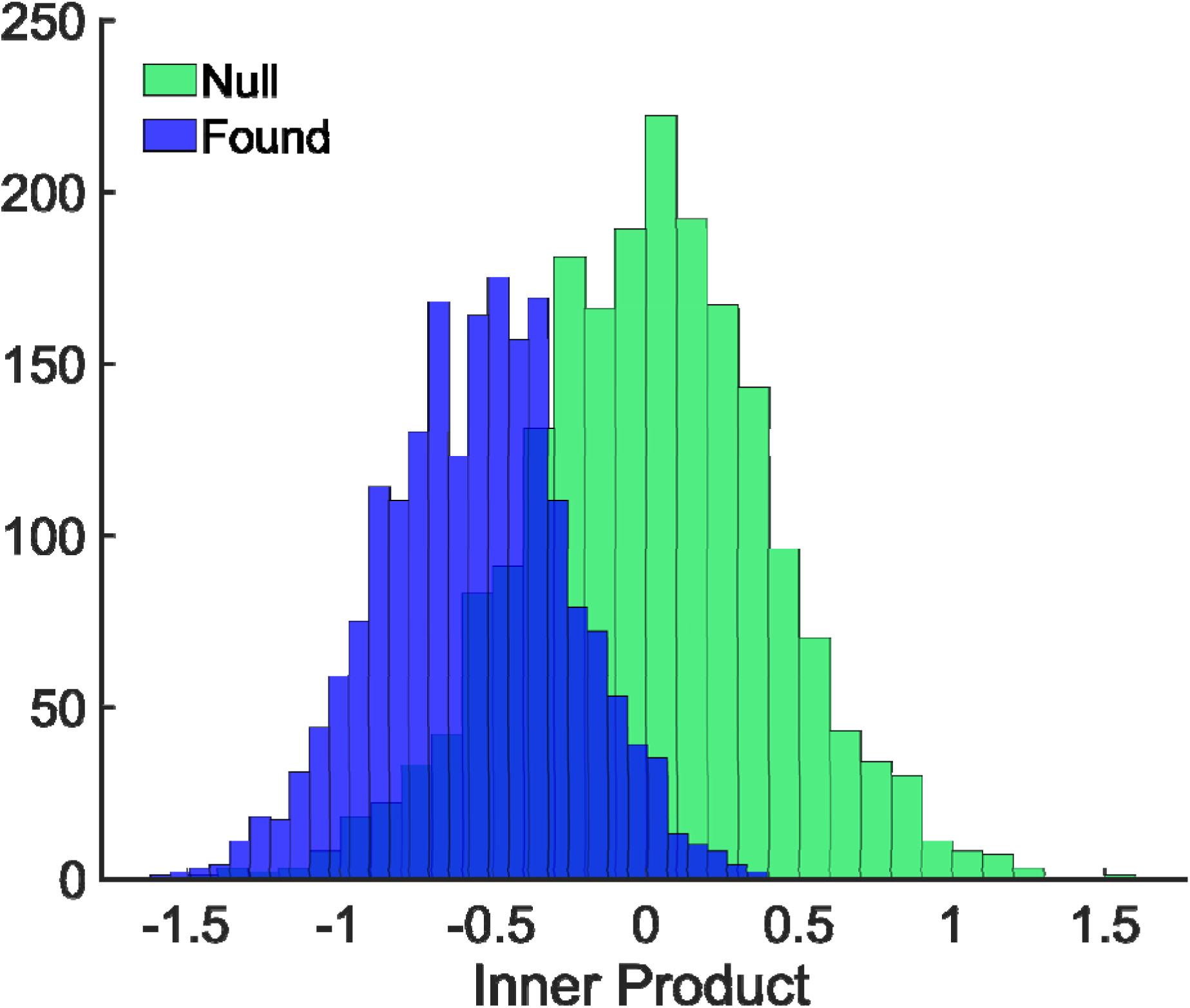
Example distribution of 2000 inner product draws from the Null and the linear model for single-trial spectra. The distribution of samples suggest the opposite pattern to what is typically observed in aversive conditioning paradigms (as evidenced by the negative-valued inner product distribution).

For single spectra methods, ssVEP findings indicated opposite trends to typical amplification of the CS+ condition in aversive conditioning experiments. Bayes factor comparisons showed decisive evidence in favor of a generalization relationship among conditions, (Log_10_ BF= −1.18) over an all-or-nothing pattern (Log_10_ BF= 0.96) in the opposite direction than what has been found aversive conditioning research.

**Figure 7.**
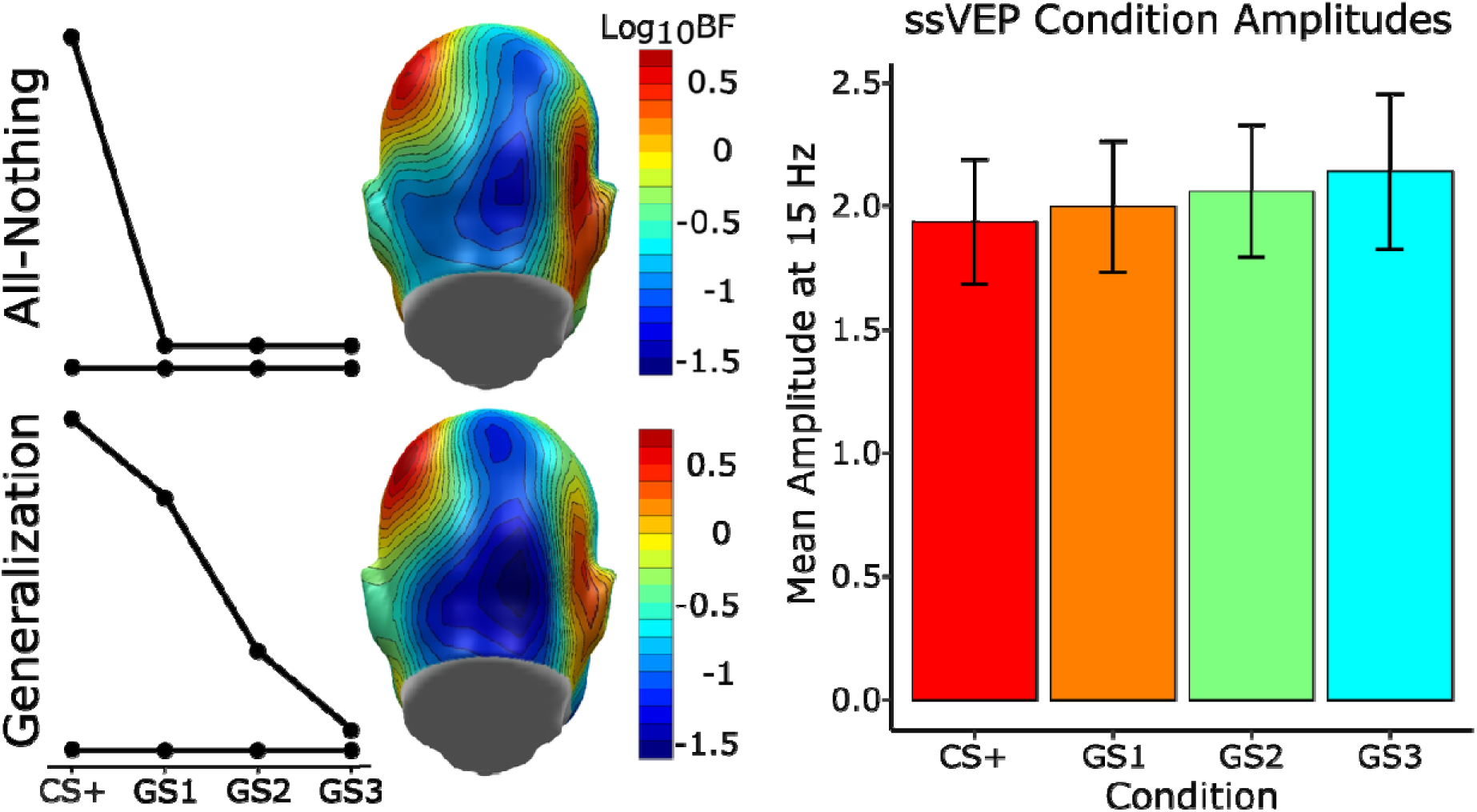
Panel A) ssVEP dot product with model weights converted to Log10 Bayes Factors. Top A: Log_10_ BF for all-nothing model. ssVEP topography shows no decisive evidence for this pattern. Bottom A: Log _10_ BF for Generalization model. ssVEP topography shows evidence for the opposite generalization pattern as compared to the null model. Panel B) Raw 15 Hz ssVEP amplitudes by condition at sensor 75 show the opposite to expected generalization pattern.

### Alpha-band Activity

Figure 8 shows the time-frequency plot for both raw amplitude and percent baseline corrected amplitude at sensor 75 (Oz). To examine alpha activity specifically, we selectively extracted the time-varying alpha amplitude from the wavelet tuned at 10.56 Hz as a time series.

**Figure 8.**
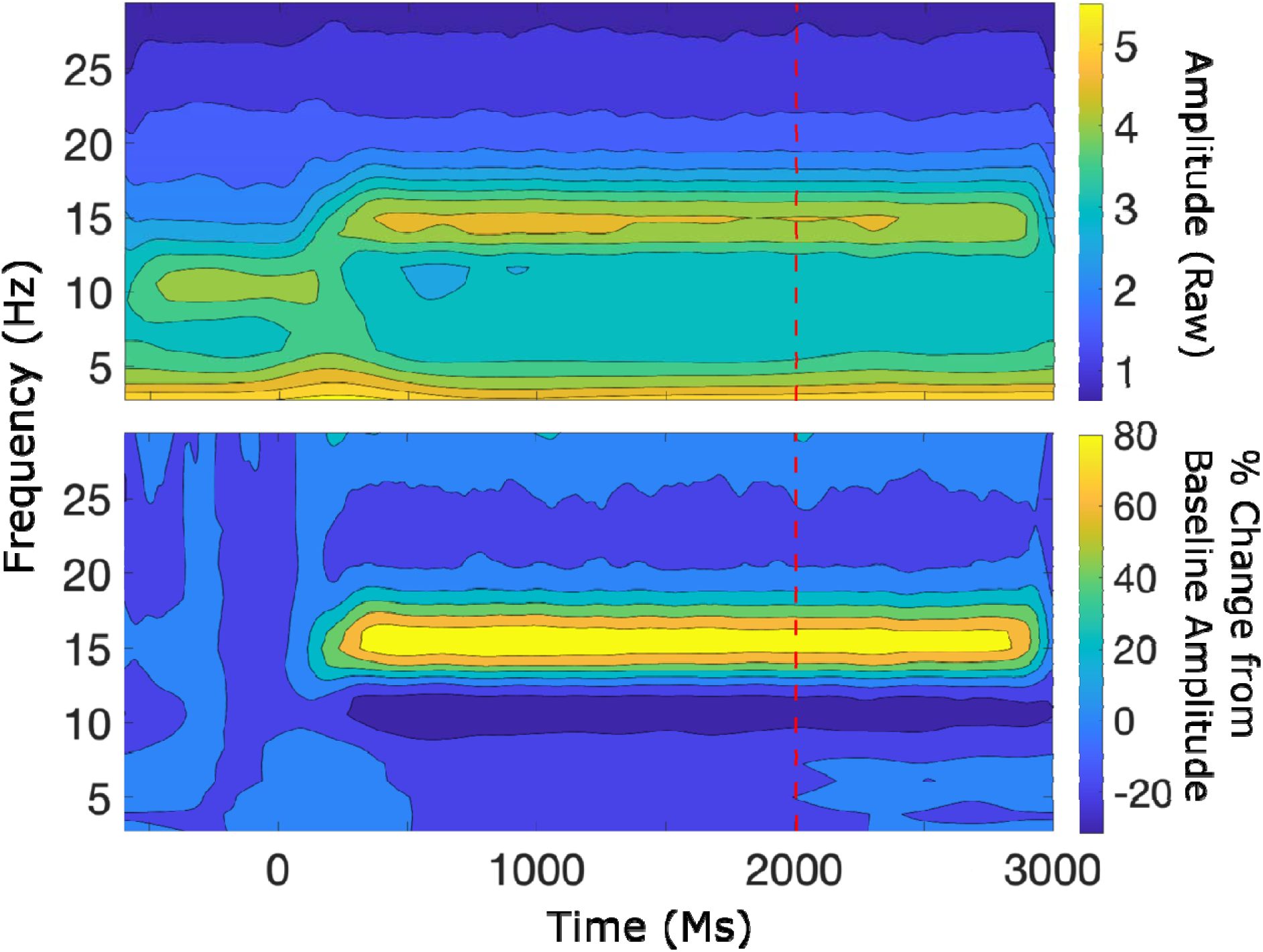
Time-frequency plot derived from EEG data at channel 75 (Oz) averaged across conditions. A) raw frequency EEG data averaged across all participants and conditions. B) Baseline corrected (percent difference) time-frequency data averaged across all participants and conditions.

Both alternative models were compared against a null model. Alpha-band activity showed no substantial evidence of fit to the generalization model in left occipital areas. A model displaying a pattern of typical alpha reduction weights of the generalization model showed no substantial effect in the occipital area during the late epoch (Oz: Generalization model, Log_10_ BF_10_ Early = −0.13, Middle = −0.36, Late = −0.28).

Support for the all-or-nothing model was found, but the direction was in the opposite direction that is typically observed for alpha activity. The model showed systematic increases across averaged early, middle, and late time periods at the left-occipital electrode (sensor 70). Negative values indicate support for the opposite model to the one tested. In the early time window (∼0-700 ms) evidence was moderate (Log_10_ BF_10_ = −0.59) in favor of the opposite model. During the middle time window (∼600-1200 ms), evidence was strong (Log_10_ BF_10_ = - 1.01), and in the late window (∼1200-1800) there was also strong evidence in support for the all- or nothing model in the opposite direction (Log_10_ BF_10_ = −1.22). See Figure 9.

**Figure 9.**
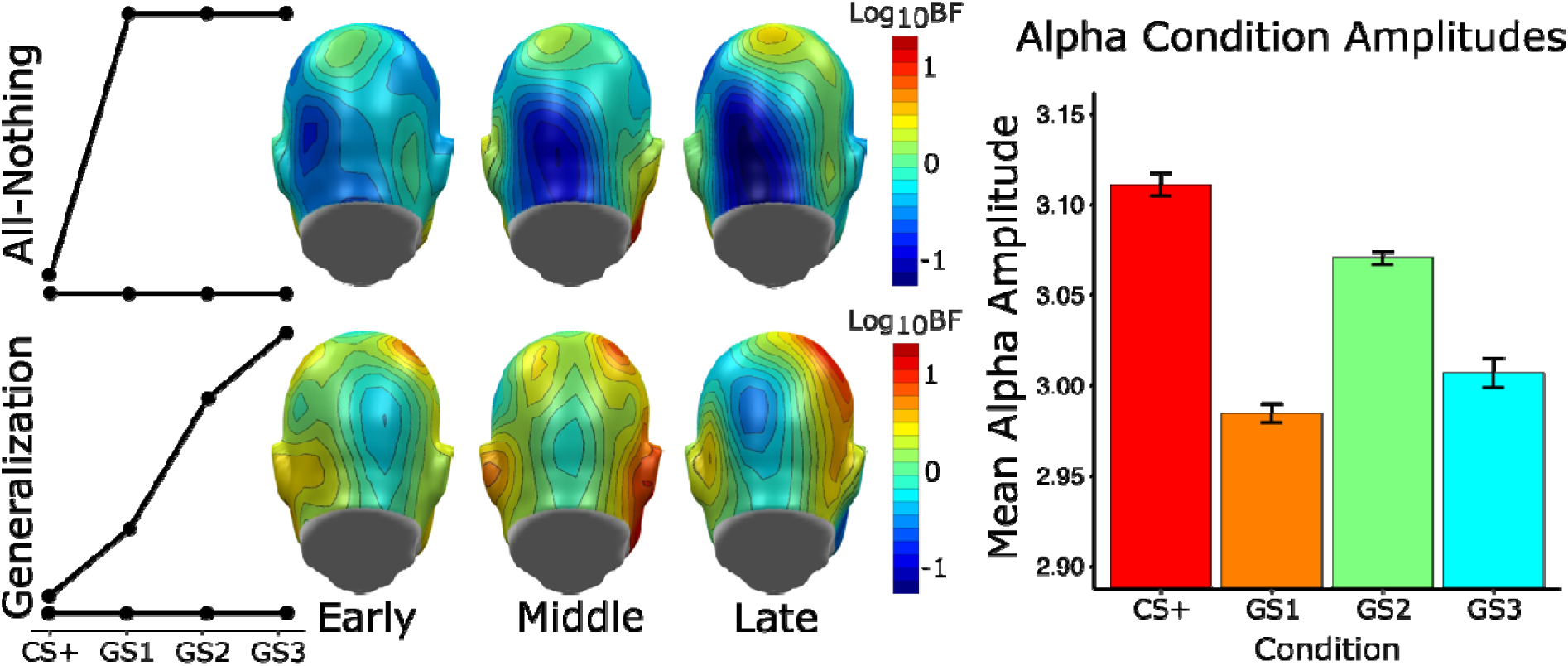
Left Panel: Transient alpha amplitude dot product with model weights converted to Log10 Bayes Factors; Early: 0-700 ms, Middle: 700:1300 ms, Late: 1300:2000 ms. Right Panel: Mean alpha amplitudes by condition averaged over 2000 ms. Typical expected model with alpha suppression for aversive conditioning in black with the found pattern in red. Top panel: All-nothing pattern and Log_10_ BF topographies for all-nothing model across early, middle, and late acquisition. Alpha topography shows decisive evidence for this pattern in early and late time bins. Bottom panel: Generalization pattern and Log _10_ BF topographies. Alpha topography shows no decisive evidence for this pattern.

## Discussion

The objective of the current study was to determine the extent to which association formation between a conditioned visual cue and a non-aversive outcome elicits similar visuocortical and oscillatory EEG responses as are observed in aversive conditioning paradigms. We utilized a paradigm identical to Ahumada et al (2024) but with a soft tone instead of a loud noise or electric shock acting as the unconditioned stimulus. Our findings regarding visuocortical activity strongly differed from the aversive conditioning literature, where increased ssVEP amplitude is consistently observed in response to threat-predictive stimuli as opposed to safety cues (McTeague et al., 2015; Stegmann et al., 2020). Previous research has demonstrated the effect of neutral and emotionally salient pictures on ssVEP responses (Keil et al., 2003). Keil et al (2003) found higher ssVEP amplitude for pleasant and unpleasant pictures as compared to neutral ones, suggesting that emotional arousal enhances visual cortical processing via re-entrant modulation.

Notably, we observed—using a pipeline that mirrored the approach used in Ahumada et al. (2024)—a pattern opposite to the former study, with the present study observing heightened ssVEP amplitudes in response to the safest generalization cue, and the lowest amplitude in response to the CS+. This finding however was restricted to the single-trial DFT analysis without the RESS approach. As such, this smaller and unexpected effect will require replication, and may reflect differences in noise level in the CS and GS conditions, eliminated by the RESS algorithm. As such, it should be interpreted with care.

SsVEP responses to aversive stimuli in conditioning paradigms are often interpreted as reflecting enhanced engagement with motivationally relevant stimuli. As such, the present study may suggest several hypotheses to be systematically examined in future work: (1) Paralleling intermodal tasks, it is possible that anticipating a neutral auditory cue in the future may reduce visuocortical engagement with a visual cue, as cross-modal attention shifts from the visual to the auditory domain (Saupe et al., 2009). (2) based on attention theories emphasizing entropy and information content of stimuli, it is also conceivable that observers engage more with GS stimuli, which are associated with higher uncertainty than the CS+ (Pearce & Hall, 1980). For example, animal model and human data suggest that attention may be directed to stimuli with higher uncertainty and to stimuli that promote learning through eliminating alternative models to account for learned contingencies (Gottlieb, 2012). The present study is unable to examine these hypotheses, but future studies may manipulate contingencies and aversiveness of CS-US pairings systematically to explore the roles of anticipation vis-à-vis processes related to entropy and information in non-aversive association formation.

Similarly to the ssVEP results – which deviate from the usual pattern – the alpha amplitude results likewise depart from consistently observed suppression of alpha-band activity in response to conditioned threat cues (Bacigalupo & Luck, 2022; Friedl & Keil, 2020; Pouliot et al., 2025): in the present Pavlovian conditioning paradigm, the CS+ elicited reduced alpha suppression. In aversive paradigms, pronounced alpha suppression is interpreted as a neural marker of vigilance to threat, facilitating enhanced sensory processing of potentially dangerous cues through increased thalamo-cortical communication and top-down attention control (Bacigalupo & Luck, 2022; Friedl & Keil, 2020; Klimesch, 2012). In contrast, the attenuated alpha suppression observed in our neutral paradigm suggests a different mechanism at work.

Specifically, attenuated alpha suppression showed patterns of all-or-nothing activity at two distinct time segments in the opposite direction to what is typically observed (see Figure 9): First in the middle time window and again in the late time window (approximately 700-2000 ms pre-tone onset). This transient activity suggests anticipatory gating of attention, where the brain modulates sensory excitability in preparation for expected events. The lack of alpha suppression, typically observed in aversive conditioning or tasks with high motivational salience, points to a reduced need for engagement in the current neutral associative learning task.

This pattern aligns with theoretical notions that alpha oscillations serve as a dynamic inhibitory mechanism, regulating sensory input based on task demands (Gratton, 2017; Klimesch, 2012). The sensory inhibitory mechanisms are observed during working memory tasks, where increased or sustained alpha amplitude is thought to gate sensory input and protect internal representations from distraction (Bonnefond & Jensen, 2012). This allows for enhanced focus on behaviorally relevant stimuli. When events and stimuli are “neutral” or lack behavioral significance, there is less need for enhanced sensory processing. Supporting this theory, Pavlov and Kotchoubey (2022) found that approximately 60% of visual working memory studies report increases in alpha-band activity across a range of stimuli and paradigms. This view is further supported by studies showing that alpha amplitude increases in anticipation of distractors and predicts better recall performance (for review see Jensen, 2024). Thus, the above hypotheses outlined for ssVEP activity apply to alpha activity as well. However, evidence for alpha activity is more abundant, as it has been examined in anticipatory, working memory, and imagining studies more frequently than ssVEP.

The results from participant ratings support this interpretation. Participants did not differ in their subjective valence judgements across conditions or over the four experimental phases, suggesting that the perceived pleasantness or unpleasantness of the stimuli remained stable throughout the course of the experiment. Arousal ratings were higher during acquisition for the CS+ compared to the GS2 and GS3. Robust main effects of both Cue and Phase on expectancy, as well as their interaction, confirm that participants formed and updated predictions about stimulus-outcome contingencies. Expectancy ratings were thus consistent with learning the contingencies during acquisition and parallel increases in alpha activity.

Another possible explanation for the transient all-or-nothing attenuated suppression, could be the absence of motivational salience in our paradigm. Previous research in aversive conditioning and emotional picture viewing has identified enhanced alpha suppression in relation to motivationally salient and emotionally valenced stimuli. For example, aversive conditioning studies typically report alpha suppression in conditions pairing neutral stimuli with aversive events (e.g., loud noise; Bacigalupo & Luck, 2022; Farkas et al., 2024; Pouliot et al., 2025). Additionally, emotionally salient pictures have been shown to elicit decreased alpha amplitude as compared to neutral pictures (for review, see Codispoti et al., 2023). From an evolutionary perspective, this reflects the idea that emotional stimuli engage neurophysiological systems to adapt behaviors in the presence of threat or reward (Lang & Bradley, 2010). In contexts where stimuli are not emotionally or motivationally significant, alpha-mediated gating may occur only transiently, sufficient to filter out irrelevant input without sustained vigilance. Furthermore, the timing of these alpha modulations may correspond to certain cognitive processes, such as expectancy updating, as suggested by studies linking alpha-band oscillations to trial-level changes in expectation (Riels et al., 2021).

Finally, the transient nature of the alpha oscillatory activity to time points right before both cue and tone suggest possible anticipatory activity. An anticipatory increase in alpha activity is observed in context where the brain prepares for an expected sensory event. This activity is thought to reflect efforts to filter out predictable input, prioritizing novel or externally generated information and preventing sensory overload (Stenner et al., 2014). This response has been observed in attention tasks, where higher anticipatory alpha is likened to reduced cortical excitability and less perceptual sensitivity (for review see Foxe & Snyder, 2011).

Several limitations should be acknowledged. Firstly, the study did not include a direct, within-person comparison to aversive outcomes, making it difficult to determine the extent to which the observed activity reflects non-aversive association formation or the broader features of the experiment, such as the nature of the stimuli or task structure. Second, the study did not incorporate additional physiological measures, such as pupil response or heart rate variability, which could provide complementary insights into participants’ autonomic responses. Third, the findings are based on a smaller sample size of undergraduate students. Future work should aim to replicate these findings using a broader range of cues and participant samples to establish the robustness of these findings. Additionally, future studies should manipulate motivational salience systematically (e.g., using positive, aversive, and neutral experiences) to better characterize how salience influences early sensory engagement. Trial-by-trial modeling of alpha dynamics and associative strength could further clarify the temporal course of learning-related neural processes for less motivationally salient association formation.

## Conclusion

Our results demonstrate that neutral associations elicit neural activity patterns that are categorically distinct from those observed in aversive conditioning paradigms. Specifically, we found less alpha suppression in the CS+ condition compared to other conditions, contrasting with the robust alpha suppression typically observed in aversive learning. Additionally, ssVEP activity patterns aligned best with the all-or-nothing model, but again, in a direction opposite to that seen with aversive stimuli at multiple time periods. Together these findings highlight that the sensitivity of both alpha amplitude and ssVEP amplitude to association formation highly depends on experimental context and the nature of the unconditioned stimulus.

